# DENSITIES, PLANT SIZES, AND SPATIAL DISTRIBUTIONS OF SIX WILD POPULATIONS OF *LOPHOPHORA WILLIAMSII* (CACTACEAE) IN TEXAS, U.S.A

**DOI:** 10.1101/2020.04.03.023515

**Authors:** Anna Ermakova, Carolyn V. Whiting, Keeper Trout, Colin Clubbe, Martin K. Terry, Norma Fowler

## Abstract

*Lophophora williamsii* (Cactaceae) is thought to be threatened by habitat loss and overharvesting. However, basic demographic and habitat information to evaluate its conservation status has been lacking. We surveyed six wild populations of this species, three in South Texas and three in West Texas, to begin to address this gap. We found high levels of heterogeneity in plant presence and density at multiple spatial scales. While plant densities were not consistently different between South and West Texas, plants were significantly larger in West Texas. The two regions differ strongly in precipitation, temperature, elevation, and topography, all of which are correlated at the regional scale. Therefore, it was not possible to identify which of these variables, or other factors such as competition and human harvesting, may be responsible for the regional differences in plant size. However, our results provide initial information for determining the conservation status of this species.

**RESUMEN:** *Lophophora williamsii (Cactaceae)* se considera amenazada por la pérdida de hábitat y cosecha excesiva. Sin embargo, se carece de información demográfica y ambiental básica para evaluar su estado de conservación. Para abordar este déficit, examinamos seis poblaciones salvajes de esta especie (tres en el sur y tres en el oeste de Texas respectivamente). Encontramos altos niveles de heterogeneidad en la presencia y densidad de plantas en múltiples escalas espaciales. Las densidades no son consistentemente diferentes entre el sur y el oeste, pero las plantas son significativamente más grandes en el oeste. Las dos regiones difieren notablemente en precipitación, temperatura, elevación y topografía. Todas estas variables están correlacionadas a escala regional, por lo que no es posible identificar cuál de ellas (u otros factores como la competencia y la cosecha humana) causan las diferencias regionales observadas en el tamaño de la planta. Nuestros resultados proporcionan información fundamental para determinar el estado de conservación de esta especie.

## INTRODUCTION

Like many other plants, many species of cacti (Cactaceae) are endangered by the degradation and loss of their habitat (Boyle & Anderson 2002; Ortega-Baes et al. 2010; Goettsch et al. 2015). Many species of cacti are also endangered by the harvesting of wild plants (Oldfield 1997). For this reason, CITES has listed the entire Cactaceae family in its Appendix II, limiting and regulating international trade in all cactus species (CITES 2019). While most of the harvest of wild cactus plants is for the horticultural and hobby markets (Robbins 2003), *Lophophora williamsii* (common names: peyote, híkuri, etc: Rojas-Aréchiga & Flores 2016) is harvested and sold for its psychoactive properties. In the United States, *L. williamsii* is legally used by the Native American Church, and its collection (with landowner permission) and sale by licensed distributors to members of this church are also legal in the United States (American Indian Religious Freedom Act 1996; Terry & Trout 2017). Otherwise its possession, sale, and use are illegal in the United States (Controlled Substances Act 1970). It is likely that past and present levels of legal and illegal harvesting of *L. williamsii* in the United States have been greater than the populations can sustain (Terry & Mauseth 2006). In this it is similar to other plants over-harvested for the actual or presumed medicinal or recreational properties of their secondary compounds, such as ginseng (*Panax quinquefolius*; McGraw et al. 2013) or *Dendrobium* species (Liu et al. 2014). *L. williamsii* has been ranked as vulnerable in the IUCN Red List (Terry 2017).

The distribution of *L. williamsii* in the United States is limited to Texas, although it also grows in several northern Mexican states (Terry et al. 2011). It may once have had a continuous range across much of southern Texas, but at present it is known to occur only in limited areas in South Texas and, separately, in West Texas (Powell et al. 2008). In South Texas, it grows on calcareous soils that support Tamaulipan thornscrub, a shrubland community dominated by species of *Prosopis*, *Senegalia* (formerly *Acacia*), and other shrub species (Jahrsdoerfer and Leslie 1988; Návar et al. 2001). In West Texas, it grows on calcareous soils that support Chihuahuan Desert vegetation (Aide & Van Auken 1985; Hernández & Bárcenas 1995; Hernández & Bárcenas 1996). Habitat loss has been particularly severe in South Texas due to oil field development, ranching (especially root-plowing to remove woody plants), and an increasing human population. As a result of habitat destruction in combination with legal and illegal harvesting of *L. williamsii*, the species is of conservation concern (Terry et al. 2011). However, the species has not been listed under the US Endangered Species Act.

In this study we report the first quantitative estimates of *L. williamsii* spatial distribution patterns, population densities, and plant sizes from six sites, three in South Texas and three in West Texas. While this information is not sufficient to evaluate the degree of endangerment of this species in Texas, this study provides important information for developing a conservation assessment of this species. Such an assessment would include an estimate of its risk of extinction as well as recommendations for conservation measures. Our results are particularly valuable because access to *L. williamsii* populations, almost all of which are on private land, is very limited.

We were also interested in the effects of environmental factors such as precipitation on this species, and therefore obtained and analyzed the relationships between a set of environmental variables and *L. williamsii* size and density. Initially, we did not have strong specific hypotheses about the effects of environmental variables, with one exception: based on past observations of one of the authors (MKT), we hypothesized that plants would be disproportionately common on south-facing slopes. Many cactus species are well-known for their adaptations to water scarcity (e.g., Nobel 2003), so one might expect West Texas to be a more favorable environment for this species. However, South Texas is better known for its *L. williamsii* populations than West Texas is (Trout & Terry 2016).

## MATERIALS AND METHODS

### Study sites

Study sites were located in both regions of Texas in which *Lophophora williamsii* occurs: the Tamaulipan thornscrub in South Texas and the Chihuahuan Desert in West Texas (Fig. 1, Table 1). Three sites were surveyed in each region. The six sites each met two criteria: a known population of *L. williamsii* and permission to survey the site. All six sites were in private ownership, and verbal consent was obtained from the landowner prior to surveying each study site.

**Table 1.** Study sites, suitable habitat, and areas surveyed.

| Site | Region | Ecoregion | County | Land use | Site area (ha) | Suitable habitat (ha) | Transects surveyed |
| --- | --- | --- | --- | --- | --- | --- | --- |
| 1 | South Texas | Tamaulipan thornscrub | Starr | ranch | 197.93 | 118.15 | 27 |
| 2 | South Texas | Tamaulipan thornscrub | Jim Hogg | conservation | 243.08 | 75.79 | 31 |
| 3 | South Texas | Tamaulipan thornscrub | Starr | conservation | 183.02 | 73.66 | 26 |
| 4 | West Texas | Chihuahuan Desert | Val Verde | ranch | 74.96 | 74.96 | 14 |
| 5 | West Texas | Chihuahuan Desert | Terrell | ranch | 64.37 | 52.06 | 18 |
| 6 | West Texas | Chihuahuan Desert | Presidio | conservation | 725.26 | 375.35 | 5 |

**Fig. 1.**
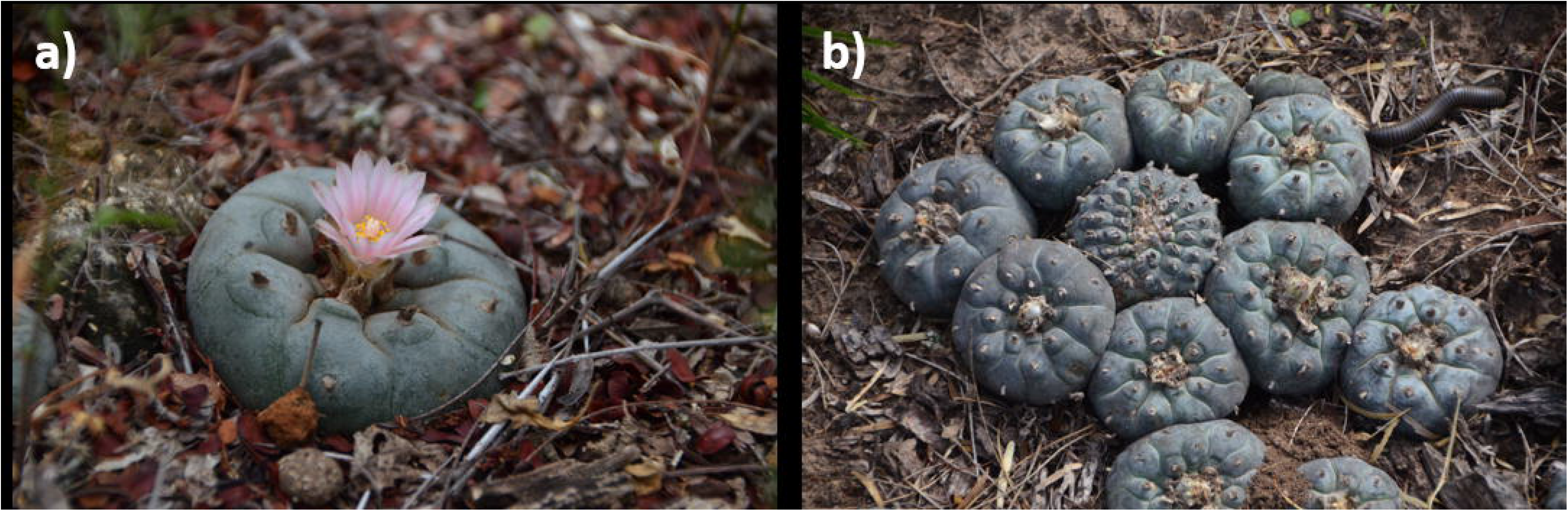
Locations of the six study sites, three in South Texas in Tamaulipan thornscrub vegetation (sites 1-3) and three in West Texas in Chihuahuan Desert vegetation (sites 4-6).

### Field study design and data collection

Site surveys were conducted in May and July 2019. Within each site, our survey methodology was designed to sample the population in an unbiased way while optimizing the inevitable trade-off between statistical rigor and sample size.

For each site, we first identified the area to be sampled in that site. The area to be sampled was defined as all of a given site that

(a) was accessible, which was defined as being within 200m of a walkable road or trail, no further than 2 km from vehicle access, and not on a dangerously steep slope; and
(b) was suitable habitat, defined as

(b1) did not appear to have ever been root-ploughed or converted to agriculture (which each create lasting changes in the vegetation); and
(b2) had not been developed (i.e., no roads, buildings, pipelines, or similar disturbances were apparent); and
(b3) had white or gray soil, which indicated that the bedrock was limestone and the soil calcareous (soil color visible in aerial photographs; soil type checked with NRCS soil maps); and
(b4) was not near a stream, dry drainage channel, or riparian vegetation.

The open-source geographic information system QGIS (v. 3.8.2; QGIS Development Team) was used to randomly place transects within the area to be sampled in each site. To avoid bias in transect location and to make transect location in the field easier, each transect was oriented north-south. These parallel transects were at least 250 m apart (as measured east-to-west, i.e., perpendicular to the transects). Each transect was 25 m long and 4 m wide. A random set of possible transects was generated in advance for each site, from which a random subset was selected while at the site; in this way we adjusted the number of transects per site to our available time and other logistic constraints at each site. We used a Garmin GPSMAP 64s GPS unit with 3-5 m accuracy to locate transects in the field (Garmin Ltd, Olathe, Kansas, USA). A total of 121 transects, divided among six sites, was surveyed (Table 1).

All individual *L. williamsii* plants in each transect were recorded. Crowns were assumed to be from the same plant if touching; if not touching, they were considered to be different plants. The number of crowns per plant and, for each crown, its shortest and longest diameter, were recorded.

### Electronic data sources

We used free, publicly available electronic data bases, accessed in 2019, to provide environmental information. The Digital Elevation Model of the Unites States Geological Survey (USGS) provided elevation, slope, and aspect; these data were supplemented by USGS topographic maps and geological maps. The Texas Natural Resources Information System (TNRIS) provided land parcel data used to determine property boundaries. The Parameter-elevation Regressions on Independent Slopes Model (PRISM) Climate Database provided 30-year average climate variables. Soil data came from the United States National Resources Conservation Service Web Soil Survey. Harvesting and sales data was obtained from the Texas Department of Public Safety. All of these variables were combined with our transect locations into a single geographically-referenced data set using QGIS.

### Calculation of plant size, plant density, and aspect

We estimated the above-ground volume of each crown as a hemisphere (i.e., having a circular base) whose basal diameter was the average of the longest and shortest measured diameters of that crown.

Above-ground volume of each crown was therefore calculated as crown volume = ⅔ π x (average diameter/2)^3^. Individuals of *L. williamsii* usually have a single crown, but some grow in caespitose clumps (Fig. 2). Multiple crowns are the result of past injury, either harvesting (Terry et al. 2011) or a natural injury such as herbivory (M. Terry, pers. obs.). If a plant had multiple crowns, the estimated volumes of each of its crowns were summed to obtain the total above-ground volume of the plant.

**Fig. 2.**
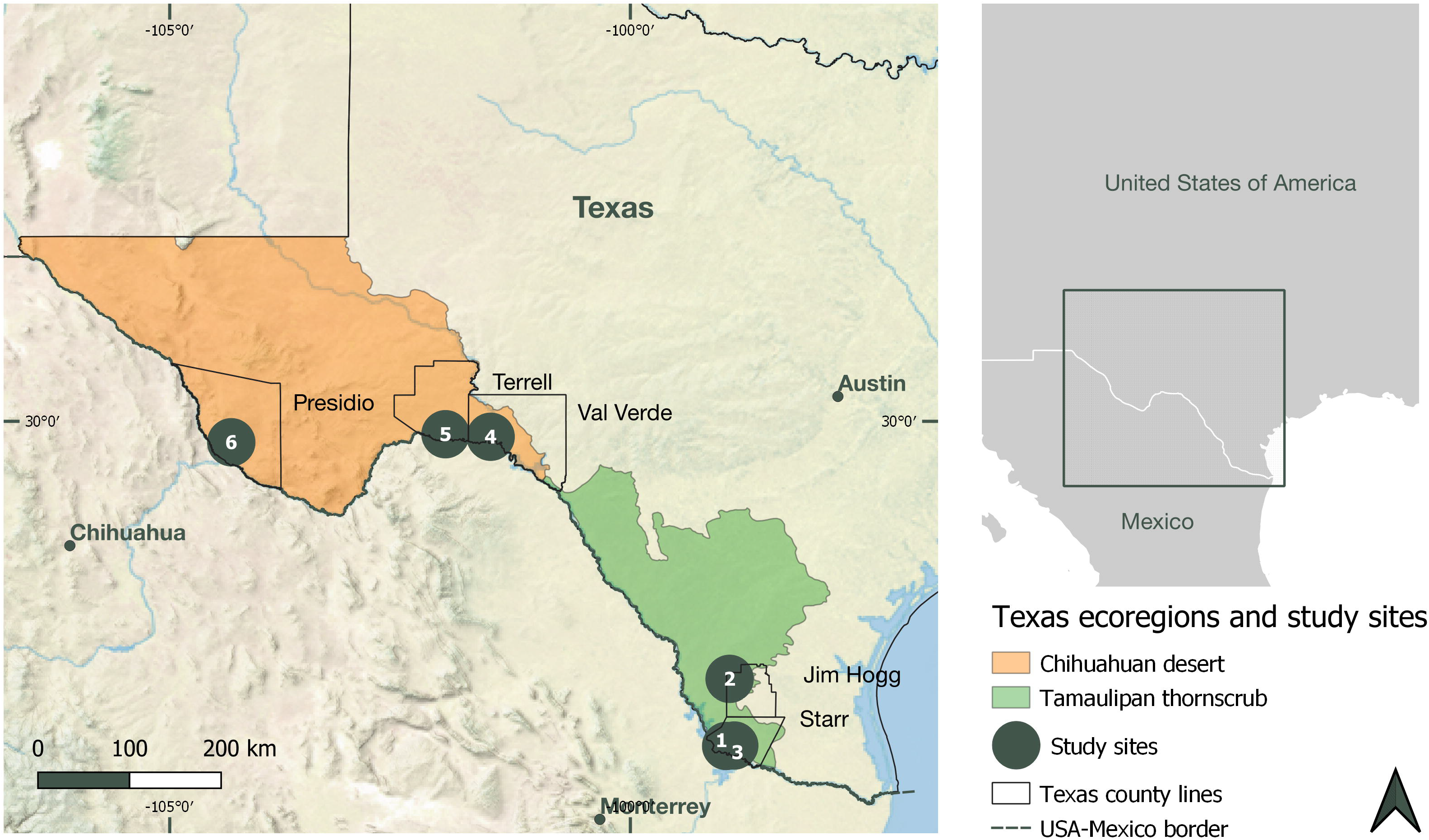
*Lophophora williamsii* (peyote) cactus (a) with a single crown, in flower, and (b) growing in multi-crown cluster. Photos by AOE.

Population density per site was estimated as the number of *L. williamsii* plants per hectares surveyed, i.e., number of plants in all the transects surveyed in that site, divided by the total area of all transects surveyed in that site.

Aspect often differed from plant to plant within a transect. Because aspect is circular rather than linear (0° = 360°), it was converted to a categorical variable (315°-45° = north-facing, 45°-135° = east-facing, 135°-225° = south-facing, 225°-315° = west-facing).

### Statistical analysis

SAS (v. 9.4, SAS Institute, Cary, North Carolina) was used for statistical analyses.

#### Variation at different spatial scales

Variation in plant volume at different spatial scales (region, site, transect, plant) was analyzed with a hierarchical analysis of variance (ANOVA), with site nested within region, transect within site, and plant (the “error” term) within transect (Table 2). Regions were compared using the site mean square as the denominator of the F-test, and sites within regions were compared using the transect mean square as the denominator of the F-test. All tests, and least square errors, were calculated using the hierarchical (Type I) sums of squares. The SAS GLM procedure, including its TEST statement, was used for this ANOVA. Normality of residuals was checked with the SAS UNIVARIATE procedure. Plant volume was log-transformed before analysis to improve normality of residuals.

**Table 2.** Sums of squares table of the analysis of spatial variation in plant size, measured as the logarithm of estimated plant volume. Nested ANOVA, hierarchical (Type 1) sums of squares. Abbreviations: df, degrees of freedom; SS, sum of squares; MS, mean square.

| term | df | SS | MS | denominator<br>of F | F | df of F | P |
| --- | --- | --- | --- | --- | --- | --- | --- |
| region | 1 | 162.18 | 162.18 | site within<br>region | 13.38 | 1,4 | 0.0216 |
| site within region | 4 | 48.47 | 12.12 | transect<br>within site | 3.19 | 4,8 | 0.0764 |
| transect within site | 8 | 30.41 | 3.80 | plant | 3.11 | 8,280 | 0.0022 |
| plant within transect<br>(i.e., 'error') | 280 | 342.72 | 1.22 |  |  |  |  |

Numbers of crowns per *L. williamsii* plant, presence or absence of *L. williamsii* plants in transects, and numbers of *L. williamsii* plants per transect (including zeros) were analyzed using generalized linear models via the SAS GLIMMIX procedure. Model fits were checked by verifying that the value of χ^2^/df of the model was close to 1. Presence/absence of *L. williamsii* in a transect was analyzed using the binomial distribution and logit link function. Numbers of plants/transect were analyzed using the negative binomial distribution and a log link function. Numbers of crowns per plant were also analyzed with a negative binomial distribution and log link function, but at the plant, not transect, level.

#### Effects of environmental variables

Except aspect, the effects of environmental variables were analyzed at the transect level. Analyses of their effects on plant size used transect mean plant size (mean of log volume; average calculated using all plants in a transect after log-transformation). Presence or absence of *L. williamsii* in each transect, and the number of plants (including zeroes) in each transect were also analyzed at the transect level. Region and site within region were included in each model (Table 3), and the effect of each environmental variable was tested using site within region as the denominator of the F-test. Precipitation, elevation, slope, and average maximum and minimum temperatures were added separately to the basic model, creating five separate models for each response variable. To account for doing multiple separate tests, the critical significance level of each test of an environmental variable was considered to be 0.0102 (Bonferroni correction, 5 tests) instead of 0.05.

**Table 3.**
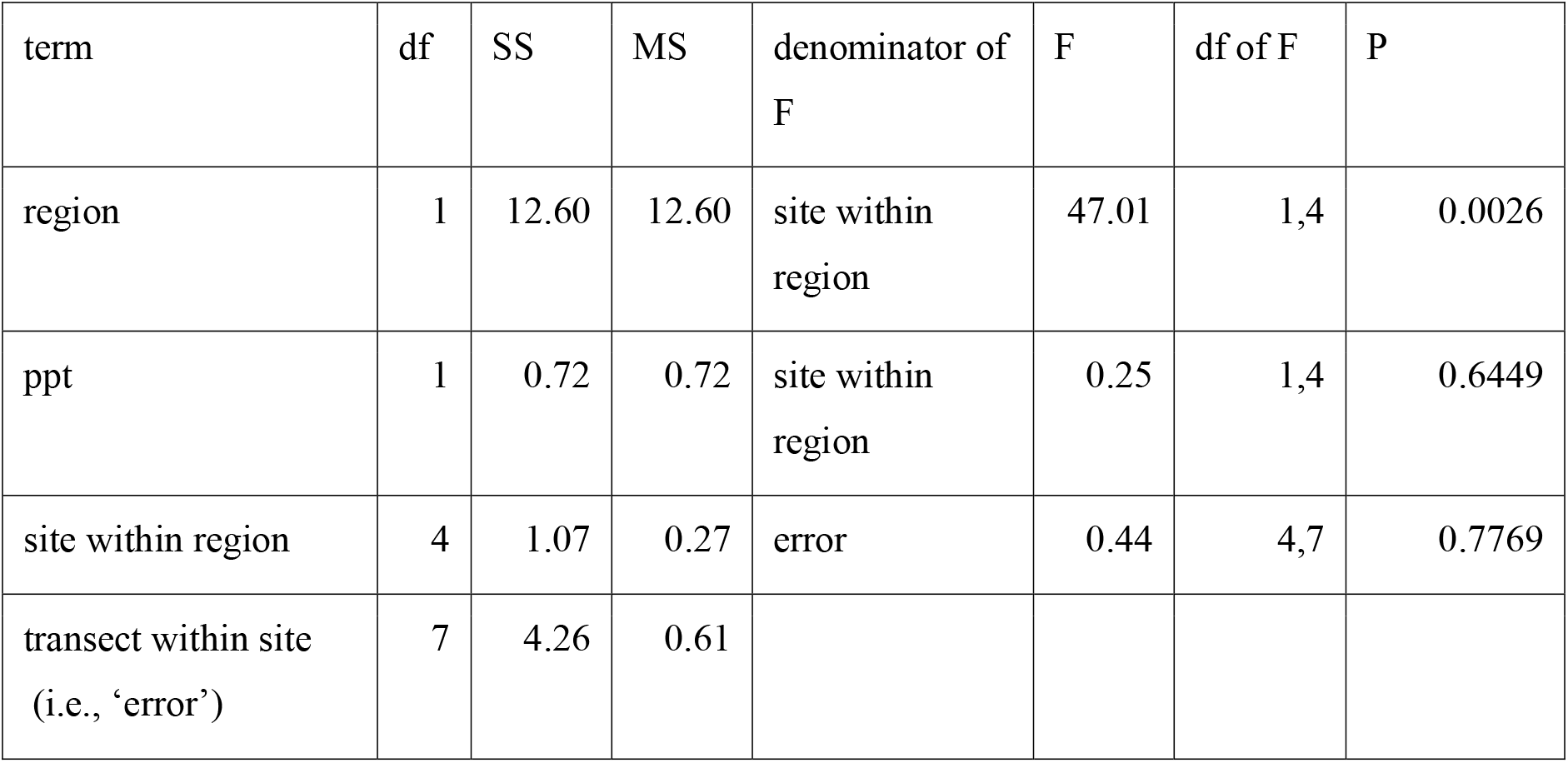
Sums of squares table of the analysis of one of the environmental variables (annual precipitation). The response variable was average (after log-transformation) of plant volume of each of the 14 transects that had at least one plant in them. Nested ANOVA, hierarchical (Type 1) sums of squares. Abbreviations: df, degrees of freedom; SS, sum of squares; MS, mean square.

| term | df | SS | MS | denominator of F | F | df of F | P |
| --- | --- | --- | --- | --- | --- | --- | --- |
| region | 1 | 12.60 | 12.60 | site within region | 47.01 | 1,4 | 0.0026 |
| ppt | 1 | 0.72 | 0.72 | site within region | 0.25 | 1,4 | 0.6449 |
| site within region | 4 | 1.07 | 0.27 | error | 0.44 | 4,7 | 0.7769 |
| transect within site<br>(i.e., 'error') | 7 | 4.26 | 0.61 |  |  |  |  |

In South Texas the slope was often so slight that aspect had little meaning (Table 4). Therefore, aspect was only analyzed for West Texas plants, where slope was always > 9%. The effects of aspect on plant volume were analyzed with an ANOVA (SAS GLM procedure) that included site and transect, plus aspect as a categorical variable (north-facing, south-facing, etc.).

**Table 4.** Average values of environmental variables (averaged across transects in a site).

| Site | Region | Average annual precipitation (mm) | Average daily maximum temperature (°C) | Average daily minimum temperature (°C) | Elevation a.s.l (m) | Slope |
| --- | --- | --- | --- | --- | --- | --- |
| 1 | South Texas | 505.81 | 30.23 | 17.07 | 88.80 | 1.60 |
| 2 | South Texas | 544.50 | 28.79 | 16.11 | 231.59 | 5.42 |
| 3 | South Texas | 504.10 | 30.15 | 17.07 | 86.48 | 1.67 |
| 4 | West Texas | 385.65 | 27.41 | 13.27 | 490.71 | 14.42 |
| 5 | West Texas | 361.22 | 27.28 | 12.89 | 532.61 | 12.92 |
| 6 | West Texas | 338.34 | 26.51 | 10.21 | 1258.80 | 13.80 |

Because we did not sample the four directions equally, we used data from Digital Elevation Model (USGS) to estimate the area of each transect that faced in each of four directions, using the definitions given above. We then, for each direction separately, summed these areas to calculate the total area we surveyed in West Texas that faced each of the four directions. These areas quantify the environment in our transects potentially available to plants. We then calculated the observed proportion of plants on south-facing slopes, and separately on west-facing slopes, each with confidence limits based on the normal approximation of the binomial and a Bonferroni correction for two tests.

## RESULTS

### Regional environmental differences

Our sites in the two regions differed substantially in elevation, slope, precipitation, and temperature (Table 4). The West Texas (Chihuahuan Desert) sites were higher in elevation, substantially drier, and colder, especially at night. The South Texas transects were close to flat, while the West Texas transects had slopes up to 25%. Because most of the variation in these variables was at the regional level, and these environmental variables were correlated with each other at the regional level, it is unclear which environmental factors were most important in determining *L. williamsii* density and size.

### Plant size

The distribution of our measure of *L. williamsii* plant size, estimated above-ground volume, had a long right tail (Supplementary Fig. 1). After log transformation of plant volume, plants were significantly (P = 0.02; Tables 2 and 5) larger in West Texas than in South Texas, 10.88 v 9.10 (least-square means of log-transformed volumes); these values are equivalent to volumes of 53.11 cm^3^ and 8.95 cm^3^ respectively, and single-crown diameters of 5.9 cm and 3.2 cm respectively. A similar pattern is seen in the raw mean volumes (no transformation, all sites and transects pooled), which were 95 cm^3^ and 22 cm^3^ in West Texas and South Texas respectively; these are equivalent to single-crown diameters of 7.1 cm and 4.4 cm respectively.

**Table 5.** Estimated plant above-ground volumes: least square means from the statistical analysis of Table 3 and their back-transformed equivalents. Some plants had more than one crown; equivalent single-crown diameters were calculated as if each plant had only one crown, and are provided only for comparative purposes.

| Site | Region | mean $\log_e$(estimated plant volume) | back-transformed plant volume (cm<sup>3</sup>) | equivalent single-crown diameter (cm) |
| --- | --- | --- | --- | --- |
| 1 | South Texas | 9.4240 | 12.38 | 3.62 |
| 2 | South Texas | 8.9158 | 7.45 | 3.05 |
| 3 | South Texas | 8.9570 | 7.76 | 3.10 |
| 4 | West Texas | 10.5332 | 37.54 | 5.23 |
| 5 | West Texas | 10.8036 | 49.20 | 5.73 |
| 6 | West Texas | 11.3039 | 81.13 | 6.77 |

The variation among sites within each region did not reach significance (P = 0.08; Tables 2 and 5). The variation among transects within sites was significant (P = 0.002; Table 2). The model explained 41% of the variation (R^2^ = 0.41), so 59% of the variation in plant size occurred among plants in the same transect.

None of the six environmental variables (annual precipitation, average minimum and maximum temperatures, elevation, slope, or aspect) explained a significant amount of the variation in plant size between sites within a region. Because the values of these six variables were so highly correlated with region, and therefore with each other at the regional level (Table 4), we cannot distinguish statistically the separate effects of these environmental variables on plant size at the regional level.

### Plant density

The six study sites (Fig 1) occupied a total area of 1489 ha, 770 ha of which were judged to be suitable habitat. We surveyed 121 transects randomly located in accessible suitable habitat (see Methods). These transects covered an area of 1.21 ha, in which we found 294 plants. The overall density of plants in suitable habitat was therefore 294/1.21 ha = 243 plants/ha. Densities differed among sites but these differences were not consistent between regions (Table 6).

**Table 6.** Plant density. All transects were placed within areas judged to have suitable habitat (see Methods).

| Site | Region | Site area (ha) | Suitable habitat (ha) | % suitable habitat | Transects surveyed | Summed transect area (ha) | No. transects with plants | No. plants on transects | Density (plant/ha) in transects | Average no. plants per transect | Average no. plants per occupied transect |
| --- | --- | --- | --- | --- | --- | --- | --- | --- | --- | --- | --- |
| 1 | South Texas | 197.93 | 118.15 | 59.7% | 27 | 0.27 | 4 | 71 | 262.96 | 2.63 | 17.75 |
| 2 | South Texas | 243.08 | 75.79 | 31.2% | 31 | 0.31 | 3 | 73 | 235.48 | 2.35 | 24.33 |
| 3 | South Texas | 183.02 | 73.66 | 40.2% | 26 | 0.26 | 1 | 53 | 203.85 | 2.04 | 53.00 |
| 4 | West Texas | 74.96 | 74.96 | 100.0% | 14 | 0.14 | 1 | 25 | 178.57 | 1.78 | 25.00 |
| 5 | West Texas | 64.37 | 52.06 | 80.9% | 18 | 0.18 | 1 | 26 | 144.44 | 1.44 | 26.00 |
| 6 | West Texas | 725.26 | 375.35 | 51.8% | 5 | 0.05 | 4 | 46 | 920.00 | 9.20 | 11.50 |

From 31% to 100% of each site was judged to be suitable habitat for *L. williamsii* (Table 6). This percentage tended to be greater in West Texas, reflecting the lower intensity of development in that region.

Plants were not equally represented in areas with different aspects. Pooling all West Texas transects, 65% (confidence limits: 54% - 76%) of those plants were growing on south-facing slopes and 35% (confidence limits: 26% - 45%) on west-facing slopes. No plants in our transects grew on east-or north-facing slopes. The area sampled by our West Texas transects was 33% south-facing, 18% west-facing, 32% north-facing, and 17% east-facing. Plants were thus significantly more common on south-facing and west-facing slopes than would be expected under a random distribution of plants. With no plants facing east or north, we cannot set confidence limits on these, but clearly plants were less common on north- and east-facing slopes than would be expected.

### Presence or absence of plants in transects

*L. williamsii* plants were highly clustered across the landscape. In South Texas 90% of transects did not have any *L. williamsii* and in West Texas 84% of transects did not (Table 6). In a model containing only region and site within region, the effect of region on presence/absence was not significant (df = 1,115; P = 0.1600), although the effect of site within region was significant (df = 4,115; P = 0.0308). None of the environmental variables added separately to this model (precipitation, minimum and maximum temperature, slope, elevation) reached the Bonferroni-corrected cutoff for significance.

### Numbers of plants per transect

As a result of the highly clustered distribution of *L. williamsii* plants, while the average number of plants per transect was less than 3 in all but one site, the number of plants per occupied transect was 12 to 53 (Table 6). Although a negative binomial distribution fitted the data well, neither region, nor site within region, nor any of the environmental variables added separately to this model (precipitation, minimum and maximum temperature, slope, elevation) were significant.

### Number of crowns per plant

Seventy-seven percent of plants had a single crown (78% in South Texas, 75% in West Texas), 14% had two crowns, and the remainder had 3-7 crowns. Numbers of crowns per plant differed significantly among sites (F(4, 288) = 4.41, P =0.0018), but not regions (F(1,288) = 1.37, P = 0.2436).

## DISCUSSION

Our results provide initial information for determining the conservation status of this species, especially estimates of density and patterns of spatial heterogeneity. We also identified regional differences in plant size and potentially important environmental factors.

### Plant size

Plant size (measured as estimated above-ground volume) in the *L. williamsii* populations we studied was log-normally distributed with a long right tail, i.e., relatively many small plants and relatively few large plants. This distribution of plant size is typical of plant populations that have more-or-less continuous recruitment; highly episodic recruitment, in contrast, often produces a distribution of plant sizes with multiple peaks (Godínez-Alvarez et al. 2003; Medel-Narvaez et al. 2006). However, even episodic recruitment can produce log-normal size distributions if the proportion of surviving plants decreases with age and there is enough variation in growth rates among plants and among years to blur the separate peaks. Therefore, we cannot rule out episodic recruitment of *L. williamsii*, perhaps related to rainfall.

There was considerable variation in plant size among plants within a transect, consistent with log-normal size distributions. In contrast, differences in plant size among sites within a region did not reach significance. Size differences between the two regions, South Texas and West Texas, were significant (see below).

Our best measure of plant size was above-ground volume estimated from crown diameters. There was also variation in the number of crowns per plant. About three-quarters of plants had only a single crown. Crown number differed significantly among sites within a region but not between regions. This latter finding was unexpected because partial harvesting, which tends to promote multiple crowns (Terry et al. 2011), is more common in South Texas.

### Spatial distribution of plants

Plants were highly clustered in space at multiple scales. Overall, only 14 of 121 randomly located transects (12%) has any *L. williamsii* plants in them. Sites differed significantly in the proportion of transects that had *L. williamsii* plants, although regions did not. A clustered distribution of *L. williamsii* was also found in the Natural Protected Area of Wirikuta, Mexico (Montero Anaya & Garcia-Rubio 2010).

Because our study was limited to three sites per region, we cannot determine the degree of clustering of plants within each of the regions. However, we expect that it is substantial, in part because this species is found only on calcareous soils, and in part because land management in South Texas (which includes root-plowing to remove woody plants in rangeland, as well as oil field infrastructure, row-cropping, and urban development) and overharvesting of this species have undoubtedly extirpated many populations.

Root-plowing, which is done to remove woody plants to improve forage availability for livestock, also kills many non-woody plant species. While some of the woody species recolonize root-plowed areas rapidly, many plant species do not (Ruthven et al. 1993).

### Plant density and species abundance

A critical component of determining whether a species is in danger of extinction is estimation of a species” abundance. We estimated plant density in our six study sites to be approximately 240 plants per hectare. However, the differences among sites in plant density reduce the utility of this value. Without site 6, an exceptionally high density site in West Texas (Table 6), the density of plants in the other five sites was approximately 214 plants per hectare, and there were only 159 plants per hectare in the two other West Texas sites.

Taken at face value, the densities we measured, when combined with our estimates of suitable habitat, produce estimates of total numbers of plants per site that are quite high. For example, we estimated that Site 1 had 118 ha of suitable habitat and a density of 263 plants per hectare, which yields an estimate of approximately 31,000 plants on this 198 ha property. However, we hesitate to put too much trust in these numbers, given the observed extreme patchiness of the spatial distribution of the species and the fact that suitability of habitat was based on aerial photographs, electronic soil maps, and similar map data, and was not ground-truthed except on the transects.

More importantly, our data cannot support an estimate of regional plant numbers because our field sites were not representative of the two regions: they were each sites already known to have *L. williamsii* present. Our sites would better be considered to represent densities characteristic of tracts of land with high conservation value for this species.

*L. williamsii* is not found in areas that have been root-plowed, and is instead associated with diverse, undisturbed plant communities (M. Terry, pers. obs.). However, past root-plowing is not always evident in aerial photographs. For this reason, and because almost all land in both regions is privately owned with no public access, we have not attempted to estimate the spatial extent of suitable habitat in either region in this study. We also do not have reliable data about where *L. williamsii* has been harvested to extirpation.

### Plant preference for south- and west-facing micro-sites

While our South Texas sites did not have enough variation in elevation to make aspect meaningful, in West Texas plants were significantly more common on west- and south-facing slopes than elsewhere. In general, we expect south-facing and west-facing microsites to be warmest, due to more hours of sunlight and sunlight during the hottest part of the day, respectively (Nobel 1978). This suggests that in West Texas the northern range limit may be set by cold temperatures, as it is for other cacti (Nobel 2003).

### Differences between South Texas and West Texas

Because our South Texas sites were consistently wetter, warmer, lower in elevation, and flatter than our West Texas sites, we cannot be sure which of these environmental variables accounts for the large difference in plant size between the two regions. All of these environmental variables are known to affect plant performance in some other cactus species (Nobel 2003). West Texas plants were larger than South Texas plants, even though our West Texas sites were drier and colder. It may be that *L. williamsii* plants experience less competition in West Texas. Consistent with this hypothesis, our casual observations are that total plant biomass of all species was much lower in our West Texas than in our South Texas sites.

However, as the southwestern United States becomes drier due to anthropogenic climate change (Ault et al. 2016), suitable habitat for this species in West Texas may become less abundant.

Other possible explanations for the differences between South Texas and West Texas *L. williamsii* plants include more harvesting and other disturbances in South Texas, resulting in younger plants there. However, the proportion of *L. williamsii* plants with only one crown was almost identical in the two regions. We also cannot rule out genetic differences between South Texas and West Texas plants.

### Implications for management and conservation

The results of our study provide a starting point for determining the conservation status of this species. We found considerable spatial heterogeneity at multiple scales in plant density. Any future conservation work should take this spatial heterogeneity into account.

If combined with more data on population dynamics (recruitment rates, survival rates, growth rates; for examples involving other cactus species, see Valverde and Zavala-Hurtado 2006; Martínez-Ávalos et al. 2007; and references therein), our results will contribute to a determination of whether harvested and/or unharvested populations are declining.

The differences we found between South Texas and West Texas are also important for the conservation of this species. At the most basic level, conclusions from one region may not apply to the other region, or two sites in Mexico. We were not able to verify the common belief that this species is much less common in West Texas than in South Texas, but neither do our results disprove it.

Our results may also contribute to a better understanding of legal and illegal harvesting of *L. williamsii*. In 2019 four registered peyote distributors operated in Texas, each employing 1 to 11 peyote harvesters (peyoteros) (Texas Department of Public Safety, https://www.dps.texas.gov). Each distributor received about 500-1500 plants per day. If we assume that the total received was 4000 plants per day, and use an estimated density of 235 plants/ha in South Texas (calculated from Table 6), 17 ha per day of suitable habitat were being harvested. Some of this harvest was likely coming from poaching on private land. Legal or not, it seems unlikely that this rate of harvest is sustainable.

## ACKNOWLEDGMENTS

Imperial College London provided funding for travel. The Cactus Conservation Institute provided equipment for AOE. We thank the landowners for kindly allowing access to their property. We also thank W. Behr, R. Carden, A. Green, T. Heinen, and B. Jones for assistance with field work, and A. Zabala Aizpuru for translating the abstract into Spanish. R. Carden and W. Behr provided comments on an earlier version of the manuscript.

## Electronic Appendix

Supplementary Fig. 1. Distributions of plant size, measured as above-ground plant volume and as the number of crowns per plant, in each study site.

