## Supplementary figures and images for "DENSITIES, PLANT SIZES, AND SPATIAL DISTRIBUTIONS OF SIX WILD POPULATIONS OF *LOPHOPHORA WILLIAMSII* (CACTACEAE) IN TEXAS, U.S.A"

### Supplementary figure 1

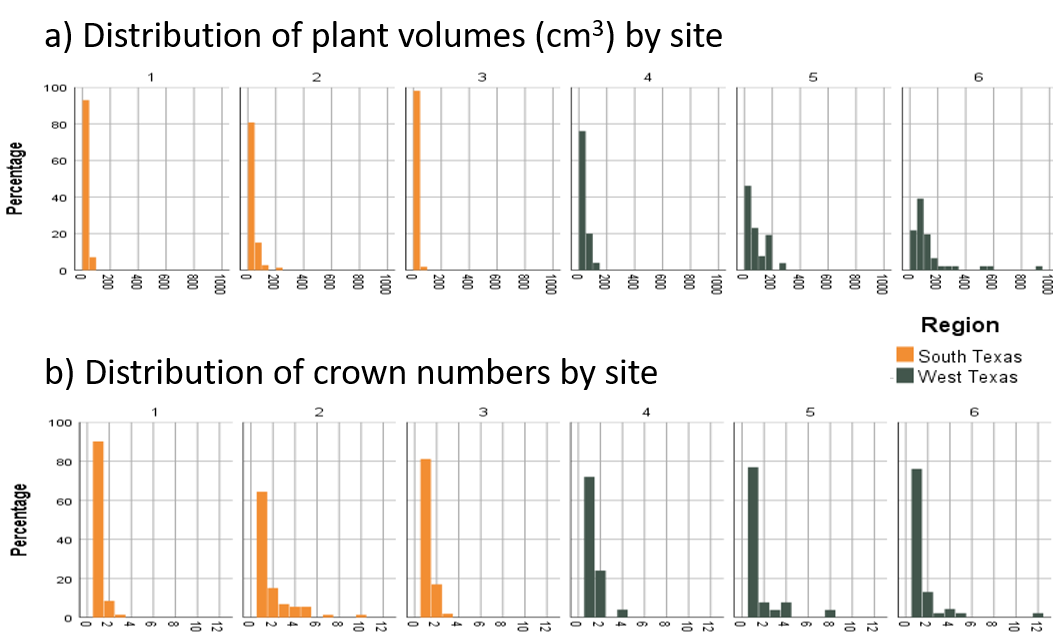
